# BAG3 regulates the specificity of the recognition of specific MAPT species by NBR1 and SQSTM1

**DOI:** 10.1101/2023.02.08.527546

**Authors:** Heng Lin, Sarah Sandkuhler, Colleen Dunlea, Darron H King, Gail V. W. Johnson

## Abstract

Autophagy receptors are essential for the recognition and clearance of specific cargos by selective autophagy, which is essential for maintaining MAPT proteostasis. Previous studies have implicated different autophagy receptors in directing distinct species of MAPT to autophagy, but the underlying mechanisms have not been fully investigated. Here we examine how the autophagy receptors NBR1 and SQSTM1 differentially engage specific forms of MAPT and facilitate their clearance. In primary neurons depletion of NBR1, unlike depletion of SQSTM1, significantly increased phosphorylated MAPT levels. The specificity of the interactions were confirmed using in vitro binding assays with purified proteins. We provide direct evidence that NBR1 preferentially binds to monomeric MAPT, while SQSTM1 interacts predominantly with oligomeric MAPT, and that the co-chaperone BAG3 regulates the specificity of these interactions. Using an in vitro pulldown assay, we show that SQSTM1 only binds to monomeric MAPT when BAG3 is absent and fails to bind when BAG3 is present. The opposite is true of NBR1; its binding to monomeric MAPT was dependent on the presence of BAG3. Interestingly, in Alzheimer’s disease brain the association of NBR1 with BAG3 was significantly decreased. In a mouse model, ablation of BAG3 in neural cells disrupted the association of NBR1 with phosphorylated MAPT and lead to increased levels of phosphorylated and oligomeric MAPT. Overall, our results uncover a novel role for BAG3 in regulating the specificity of selective autophagy receptors in targeting different species of MAPT and provide compelling evidence that BAG3 plays a key role in maintaining MAPT proteostasis.

**Highlights:**

1. First direct evidence of the district role of NBR1 and SQSTM1 in binding with monomeric and oligomeric MAPT, respectively.
2. Demonstration of a novel mechanism by which BAG3 regulates the specificity of the recognition of monomeric MAPT by NBR1 and oligomeric MAPT by SQSTM1.
3. Conditional knockout of BAG3 in the brain disrupted the association of NBR1 with phosphorylated MAPT and lead to increased levels of phosphorylated and oligomeric MAPT.

## Introduction

MAPT is a multifunctional, microtubule-associated protein mainly expressed in neurons, where it plays a key role in modulating the dynamics of microtubules [1], among other functions [2]. In pathological conditions, MAPT becomes aberrantly phosphorylated, eventually forming insoluble neurofibrillary tangles (NFTs) in a stepwise manner [3]. It has been suggested that the MAPT monomers initially interact with each other to form soluble oligomers that subsequently enlarge and precipitate as granular MAPT oligomers with a β-sheet structure which subsequently bind to each other and form insoluble NFTs [4]. The presence of NFTs is a defining hallmark of Alzheimer’s disease (AD) and other tauopathies. However, studies suggest that soluble oligomers of MAPT are likely the more toxic and pathologically significant forms of MAPT aggregates, not NFTs [5,6].

Autophagy-mediated MAPT clearance is essential for maintaining the health of the neuron and has been considered as a potential therapeutic target for AD [7]. The specificity of autophagy in recognition and selective clearance of different cargos is mediated by a family of autophagy receptors including SQSTM1, optineurin, NBR1 and CALCOCO2 (NDP52), as well as others [8]. Among these autophagy receptors both NBR1 and SQSTM1 share a conserved domain architecture, with an N-terminal PB1 domain, a ZZ-type zinc finger and a C-terminal UBA-domain [9]. However, the target cargoes of SQSTM1 and NBR1 are different. Previous studies showed that SQSTM1 colocalizes with misfolded and aggregated MAPT species and predominantly promotes the degradation of insoluble, instead of soluble, mutant MAPT [10]. Interestingly, there is data to suggest that NBR1 is able to cooperate with endosomal sorting complexes required for transport (ESCRTs) to mediate the targeting of soluble cargos for degradation [11]. These and other studies indicate that SQSTM1 and NBR1 likely may play different roles in targeting cargos, including MAPT, to autophagy to maintain proteostasis.

BAG3 is a multidomain co-chaperone that regulates autophagy in multiple cell types [12,13]. It plays an essential role in maintaining proteostasis and serves a protective role against neurodegeneration [14–16]. Previously, we demonstrated that BAG3 plays an important role in regulating MAPT clearance through autophagy [17,18], and, more recently, we found that BAG3 also regulates ESCRT-mediated MAPT clearance [19]. These findings point to the importance of BAG3 in both autophagy and ESCRT-mediated pathways. Interestingly, in HeLa cells, BAG3 was reported to associate with SQSTM1 in response to stress [20]. Nonetheless, there is still a gap in our knowledge about how BAG3 regulates selective MAPT clearance in neurons by interacting with different autophagy receptors.

In the present study, we found that BAG3 can bind both SQSTM1 and NBR1 in neurons. However, depletion of NBR1, but not SQSTM1, resulted in increased phosphorylated MAPT levels in primary cortical neurons. Interestingly, the colocalization of BAG3 with NBR1 is significantly decreased in human AD brains compared with age-matched controls. Using an *in vitro* protein binding assay, we found that NBR1 binding with purified MAPT only occurs in the presence of BAG3, while SQSTM1 binding with MAPT only occurs in the absence of BAG3. Further, we found that BAG3 promotes SQSTM1 binding with oligomeric MAPT and inhibits its binding with monomeric MAPT. Conversely, BAG3 promotes NBR1 binding with the monomeric form of MAPT, and this binding is disrupted in absence of BAG3. Additionally, conditional knockout of BAG3 in the brain disrupted the association of NBR1 with phosphorylated MAPT and lead to increased levels of phosphorylated MAPT. These findings suggest that BAG3 acts to direct the autophagy receptor to the appropriate cargo for degradation. Overall, our findings reveal a new role of BAG3 in regulating autophagy receptor and cargo interactions and provide a potential therapeutic target for enhancing pathological MAPT clearance.

## Materials and Methods

### Reagents

Constructs: lentiviral vectors: shBAG3 (5’-AAGGTTCAGACCATCTTGGAA-3’), scrRNA (5’- CAGTCGCGTTTGCGACTGG-3’) in FG12 (with an H1 promoter) [21], or pHUUG with a U6 promoter (a generous gift of from Dr. C.Pröschel, University of Rochester) with and without GFP were used [19]. shSQSTM1 (5’- GAG GAA CTG ACA ATG GCC ATG TCC T −3’) was prepared in FG12, and shNBR1 (5’- GCA TGA CAG CCC TTT AAT AGA-3’) was prepared in pHUUG. GFP-NBR1 was purchased from Addgene (#38283). Myc-SQSTM1 was a gift from T. Johansen, University of Tromso. MAPT plasmid (0N4R) was described previously [22]. FLAG-NBR1 was generated by clone the NBR1 sequence from GFP-NBR1 into pCMV-Tag 2C vector (Agilent Technologies, #211172). pGEX6P was purchased from Millipore Sigma. pT7-htau40, vector for generating recombinant MAPT, was a gift from Dr. Jeff Kuret (Ohio State University).

Antibodies: rabbit antibodies include: BAG3 (Proteintech, 10599-1-AP), MAPT (Dako, A0024), phospho-MAPT (Thr231, AT180 Thermo Fisher OPA-03156), Normal Rabbit IgG (EMD Millipore 12-370), T22 (an oligomeric MAPT antibody) (Millipore Sigma, ABN454-I). Mouse antibodies include: Myc Tag (Cell Signaling Technology, #2276), FLAG Tag (Cell Signaling Technology, 8146), TOC1 antibody (a gift from N. Kanaan, Michigan State University)[23], phospho-MAPT (Ser262) (12E8, a gift from Dr. P. Dolan), phospho-MAPT (Ser396/404) (PHF1, a gift from Dr. P. Davies), GAPDH (Invitrogen, AM4300), Normal mouse IgG (EMD Millipore 12-371). Secondary antibodies include Alexa Fluor 594 donkey-anti-rabbit, Alexa Fluor 594 donkey-anti-mouse, Alexa Fluor 488 donkey-anti-mouse, Alexa Fluor 488 donkey-anti-rabbit, and Alexa Fluor 647 donkey-anti-mouse (Thermo Fisher Scientific). Conformation specific mouse anti-rabbit IgG (Cell Signaling Technology, #5127S). Anti-rabbit IgG HRP-conjugated Antibody (Bio-rad 51962504), Anti-mouse IgG HRP-conjugated Antibody (Bio-rad 5178-2504). Biotin conjugated BAG3 antibody was generated by biotinylation of the BAG3 antibody (Proteintech, 10599-1-AP) using EZ-Link^™^ Sulfo-NHS-LC-Biotinylation Kit (Thermo Fisher Scientific, 21435).

### Animals

All mice and rats were maintained on a 12 h light/dark cycle with food and water available ad libitum. All procedures were approved by University Committee on Animal Research of the University of Rochester. BAG3^Fx/Fx^ mice (C57BL/6N-Bag3 tm1c(EUCOMM)Hmgu/H) were obtained from UKRI-MRC Harwell Institute, United Kingdom/ International Knockout Mouse Consortium. Nestin-Cre mice (#:003771) were purchased from JAX ^®^ Mice and Services. Nestin-Cre; BAG3^Fx/Fx^ (BAG3^Nes^) mice were generated by breeding Nestin-Cre; BAG3^Fx/+^ with BAG3^Fx/Fx^. The genotype type of the mice was confirmed by PCR.

All procedures were approved and performed in compliance with the University of Rochester guidelines for the care and use of laboratory animals. Both male and female mice were collected at 22–24 months of age. The tissue preparation was as previously described [24]. The samples used in Figure 2A and 2B are from a 15-month-old mice injected with scrambled AAV as previous described [24]. Briefly, mice were anesthetized and perfused with phosphate-buffered saline (PBS). The cortex and hippocampus of a half hemisphere were dissected out for immunoblot analyses and the other half for was fixed with 4% paraformaldehyde for sectioning and immunohistochemical analysis.

**Figure 1.**
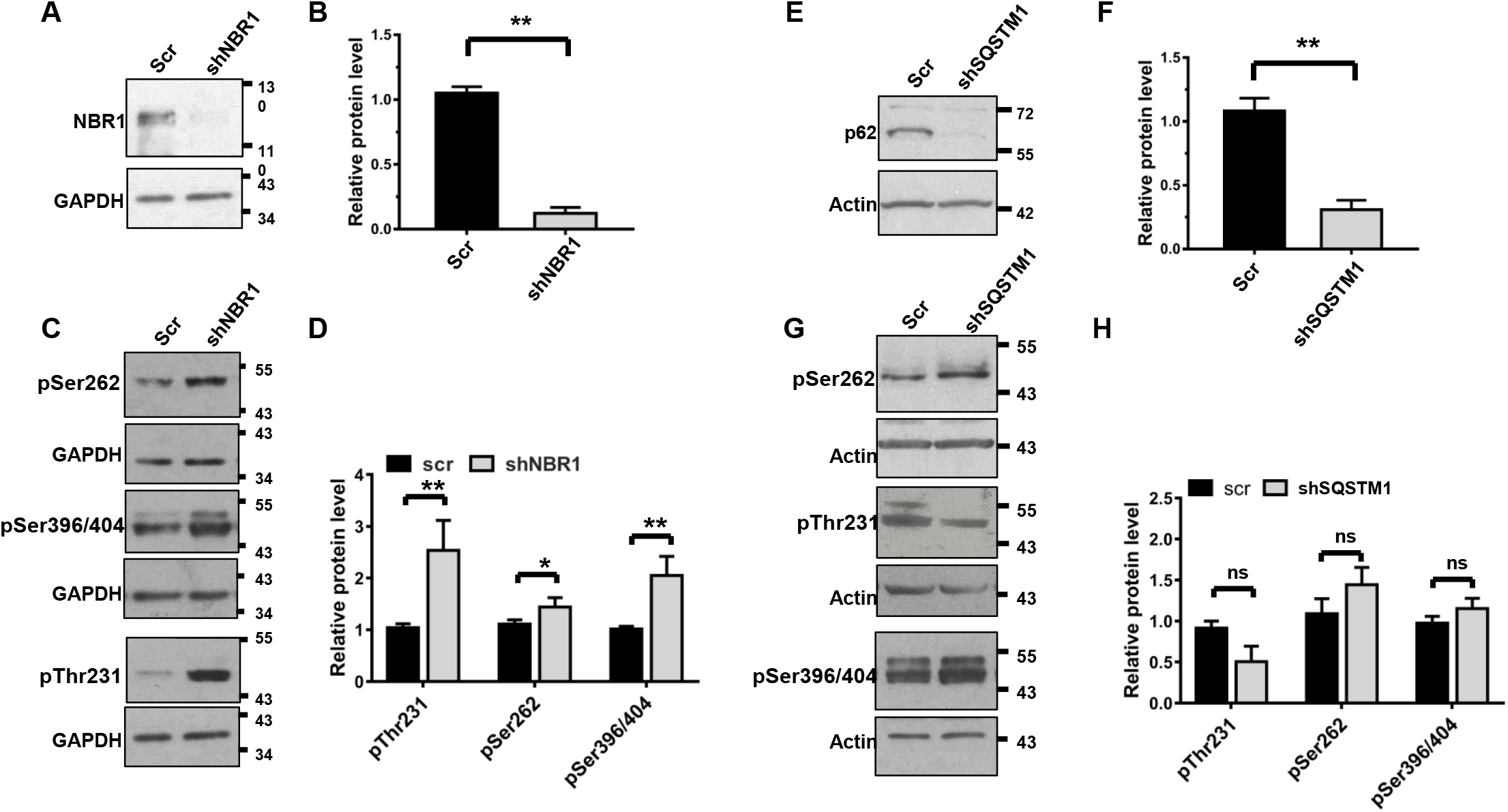
SQSTM1 and NBR1 play different roles in regulating the clearance of MAPT in neurons. (A-D) Rat cortical neurons were transduced with lentivirus expressing Scr short hairpin RNA or shNBR1. Cell lysates were immunoblotted for (**A**) NBR1, (**E**) SQSTM1 or (**C, G**) phosphorylated MAPT (p-Ser262, p-Ser396/404, and p-Thr231). GAPDH was used as a loading control. (**B, F, D, H**) Quantitation of the levels of NBR1, SQSTM1, or phosphorylated MAPT in Scr, NBR1 or SQSTM1 knockdown neurons from 3 independent experiments. Data were normalized to the loading control GAPDH or actin and then compared to scrambled controls. Data are shown as mean ± SEM. Statistical analysis was performed using the unpaired Student’s t test. *, p < 0.05; **, p<0.01.

**Figure 2.**
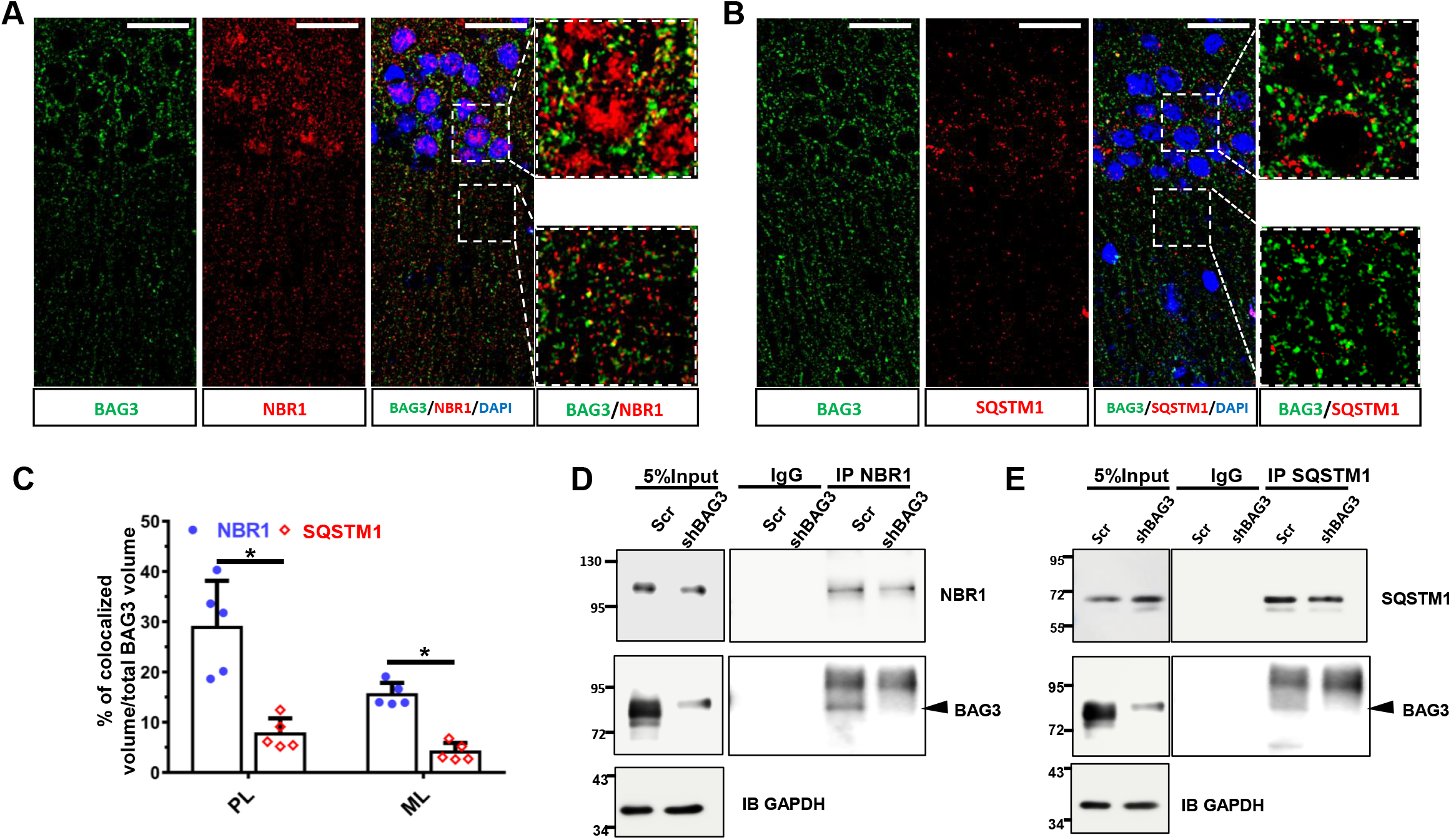
BAG3 associates with NBR1 in neurons. C57Bl/6 mice at 15-month-old were sacrificed and the brain slices were used for immunostaining analysis. (**A and B**) Fluorescence images showed the colocalization of BAG3 with NBR1(A) or with SQSTM1(**B**) in mouse CA1 region of hippocampus. Scale bars, 20μm. (**C**) Graph shows the percentage of NBR1/SQSTM1 colocalized with BAG3 volume in total BAG3 positive volume and five mice in each group were used for study. Data were plotted as mean ± SEM. *, P<0.01. PL, Pyramidal layer; ML molecular layer. (**D**) Primary cortical neurons transduced with lentivirus expressing shBAG3 or a scrambled (Scr) version. The lysate of neurons was immunoprecipitated with a NBR1 antibody and immunoblotted for BAG3 to examine the association of BAG3 with NBR1. Black arrowhead points to the BAG3 band. (**E**) The lysate of transduced neurons was immunoprecipitated with a SQSTM1 antibody and blot for BAG3 to examine the association of BAG3 with SQSTM1. GAPDH was used as loading control.

### Human brain samples

Human brain frontal cortex tissue of 6 individuals diagnosed with AD and 6 age-matched controls were obtained from the University of Florida Brain Bank. The average age of the individuals was 84.1 years for the AD patients and 82.3 years for aged controls. The aged controls were reported to be cognitively normal. Sex, post-mortem interval (PMI), Consortium to Establish a Registry for Alzheimer’s Disease (CERAD) and Thal score, Braak stage, for each case is documented (Supplementary Table 1). Brain tissues were paraffin embedded and sectioned at 5 μm.

Sections were deparaffinized with Histo-Clear II (Electron Microscopy Sciences) followed by rehydrate with gradient ethanol. Antigen retrieval with sodium citrate buffer was performed. The neurons were then permeabilized followed by blocking with PBS containing 5% BSA and 0.3 M glycine. Primary antibodies and Alexa Fluor 488/594/647 conjugated secondary antibodies were used to detect target proteins. Hoechst 33342 was used for visualization of nuclei.

### Cell culture

Primary cortical neurons were prepared from rat embryos at E18 and cultured as previously described [19]. For lentiviral transduction, DIV7 neurons were treated with virus for 16-24 hours followed by half medium change. HEK293T cells or BAG3 null HEK293T cells [19] were grown in DMEM medium supplemented with 10% fetal bovine serum (FBS), 2 mM glutamine and penicillin/ streptomycin on 60 mm dishes until 80% confluent. HEK cells were transfected with 2-3 μg of the designated construct or empty vector using the PolyJet (SignaGen Laboratories, SL100688).

### Lentivirus

Lentiviral particles were generated as previously described [17]. Briefly, lentiviral vectors were co-transfected with viral packaging vectors (pPAX (Addgene#12259) and VSV-G (Addgene#8454)) into 70% confluent HEK293TN cells using PolyJet (SignaGen Laboratories). Media was changed into DMEM containing 1% FetalClone-II serum (Hyclone, SH30066.03). Virus media was collected at 64h after transfection and filtered through a 0.2 μm syringe filter. Virus was concentrated by ultracentrifugation at 30,000 x g for 3h at 4°C, and resuspended in Neurobasal media. Concentrated virus was aliquoted, snap frozen and stored at −80°C.

### Immunoblotting

The immunoblot procedures were as previous described [19]. Cell or tissue lysates were denatured in 1x SDS sample buffer at 100°C for 10 min before being loaded onto each lane of 10% SDS-PAGE gels. After electrophoresis, proteins were transferred onto nitrocellulose membranes. Membranes were then blocked in TBST (0.1% Tween-20) containing 5% non-fat dry milk for 1h at room temperature. Primary antibodies were diluted in blocking solutions followed by incubation at 4°C overnight. The next day, membranes were further incubated with secondary antibody for 1h at room temperature. After thoroughly washing, membranes were visualized by enhanced chemiluminescence, and images captured using the KwikQuant^™^ Imager (Kindle Biosciences, LLC). The intensity of each band was quantified using Image Studio Lite (Li-Cor). GAPDH was used as loading control. Treatments were then normalized to their corresponding control sample and expressed as a fold difference above the control.

### Immunohistochemical staining of mouse brain

Brain slices (30 μm) were prepared as previously described [19]. Sections were mounted on poly-D-lysine coated slides (Thermo Fisher), antigen retrieval with sodium citrate buffer was performed, except when staining with the TOC1 antibody, and the slides were blocked with PBS containing 5% BSA and 0.1% tween 20 for 1 h at room temperature. The sections were incubated with primary antibody in 5% BSA in PBS overnight at 4°C. The next day, slices were incubated for 1 h at room temperature with Alexa Fluor^™^-conjugated secondary antibody including Alexa Fluor^™^ 594 donkey-anti-rabbit, Alexa Fluor^™^ 488 donkey-anti-rabbit, or Alexa Fluor^™^ 647 donkey-anti-rabbit (Thermo Fisher Scientific). Alternatively, they were labeled using the MOM kit (BMK-2202, Vector laboratories), followed by three washes with PBS and labeling with Streptavidin Alexa Fluor^™^ 488 or 647 (Thermo Fisher Scientific). The brain sections were coverslipped with ProLong Diamond Antifade Mountant (Thermo Fisher Scientific, P36961). The slides were imaged using a Nikon A1R HD laser scanning confocal microscope and recorded by NIS-Elements (Version 5.11) software. Resulting images were pseudocolored for illustration purposes.

### Immunofluorescence

Immunofluorescence was performed as previously described [19]. Briefly, neurons grown on coverslips were fixed in PBS containing 4% paraformaldehyde and 4% sucrose for 5 minutes at room temperature. The neurons were then permeabilized followed by blocking with PBS containing 5% BSA and 0.3 M glycine. Primary antibodies and Alexa Fluor 488/594/647 conjugated secondary antibodies were used to detect target proteins. Hoechst 33342 was used for visualization of nuclei. Images were acquired on a Nikon A1R HD scanning confocal microscope. Resulting images were pseudocolored for illustration purposes.

### Immunoprecipitation

Cells were lysed in ice-cold lysis buffer (50 mM Tris, 150 mM NaCl, 0.4% NP-40, 1 mM EDTA, 1 mM EGTA, pH7.4) supplemented with protease and phosphatase inhibitors. Cell lysates were briefly sonicated then centrifuged at 16,100 g for 10 minutes at 4°C. Cleared supernatants (500 μg) was mixed with 2 μg normal rabbit/mouse IgG control or primary antibody, followed by incubation at 4°C for 24h. Antibody/antigen mixture was incubated with Dynabeads M-280 sheep anti-rabbit or mouse IgG (Thermo Fisher Scientific) for 6h at 4°C in the following day. An aliquot of protein lysate was saved for input control. After thoroughly washing the beads, bound fractions were eluted in sample loading buffer by boiling at 100°C for 10 minutes. Proteins were then resolved by SDS-PAGE and immunoblotted.

### Recombinant protein preparation

The preparation of His-MAPT was carried out as previously described with minor adjustments [25]. His-MAPT and GST was prepared by transforming T7 Express Competent E. coli (#C2566I, NEB) with pT7-htau40, or pGEX6P inducing with IPTG, followed by resuspension in lysis buffer. To purify His-MAPT protein, resuspended cell pellet was immediately placed in boiling water for 5–20 min with stirring. The cell lysate was placed on ice for 10 min and centrifuged at 13,000× g for 20 min at 4°C to remove cell debris and denatured proteins. This supernatant was passed through a 5-μm filter followed by concentrated using 10K MWCO concentrator (Amicon). The concentrated lysate was fractionated using size exclusion chromatography. Elution was performed with 20 mM PBS buffer, and fractions were analyzed by SDS-PAGE. All the fractions exhibiting a single band near expected molecular weight (50-55 kDa) of MAPT were combined, concentrated using a centrifugal filter as above and stored at −80°C. The GST protein was purified as previous described [19]. Briefly, the lysate was purified using glutathione S-transferase beads (Ge healthcare, #17-5132). The proteins were eluted and concentrated with Amicon Centrifugal Filter Units 10K (Millipore) to remove excessive glutathione.

### Protein pull-down assays

#### FLAG-NBR1 pull-down assay

To make the FLAG-NBR1 prey, BAG3 null HEK cells were transfected with FLAG-NBR1 and cell lysate was collected 36 hours post transfection. Magnetic FLAG beads (MBL, #M185-11) were blocked with 5% BSA for 2 hours, followed by incubation with 500 μg cell lysate or BSA (per 25 μl beads) for 6 hours. The precipitate was washed with high salt wash buffer 7 times followed 2 times with PBS. The immobilized FLAG-NBR1 prey were split into equal fractions immediately for the pull-down assays.

To examine the prey binding with His-MAPT, the immobilized FLAG-NBR1 was incubated with 1.5 μg His-MAPT and 1 μg GST or GST-BAG3 (Abnova H00009531-P01) in the BAG buffer (25 mM HEPES, 5 mM MgCl_2_, and 150 mM KCl (pH 7.5)) containing 300 mM imidazole) for 2 hours [26]. The precipitated samples were denatured in 1x SDS sample buffer at 100°C for 10 min and analyzed by immunoblotting. To examine the prey binding with MAPT pre-formed fibrils, the immobilized FLAG-NBR1 is incubated with 5 μg MAPT pre-formed fibrils (StressMarq Biosciences, SPR-480) and 1 μg GST or GST-BAG3 (Abnova) in the BAG buffer for 2 hours. The precipitated sample were eluted with 100 μg/ml 3XFLAG peptide (Sigma-Aldrich, F4799). The eluted samples were split in half for reducing (mix with 5x SDS sample buffer at 100°C for 10 min) and non-reducing condition (mix with 2X non-reducing sample buffer (126 mM Tris/HCl (pH 6.8), 20 % glycerol and 0.02 % bromophenol blue) conditions.

#### Myc-SQSTM1 pull-down assay

The Myc-SQSTM1 pull-down assay was essentially the same as the FLAG pull-down assay with minor modifications. BAG3 null HEK cells were transfected with Myc-SQSTM1, and magnetic Myc beads were used for precipitation. Myc peptide (Thermo Scientific, 15265673) 500 μg/ml was used for to elute the precipitated samples.

### Image analysis

Images were opened and processed using Imaris ver.9.0 (Oxford Instruments). Colocalization analysis was done using Imaris. Specifically, regions of interest were selected at the molecular layer in CA1 region of hippocampus (Figures 2(A-B), or cortex (Figures 5(C), 5(F) and 6(A)). Co-localization module was used, and different fluorescence channels were thresholded to generate a new colocalization channel. Mander’s colocalization coefficients were used to quantify the overlapping of fluorescence intensity[27].

**Figure 3.**
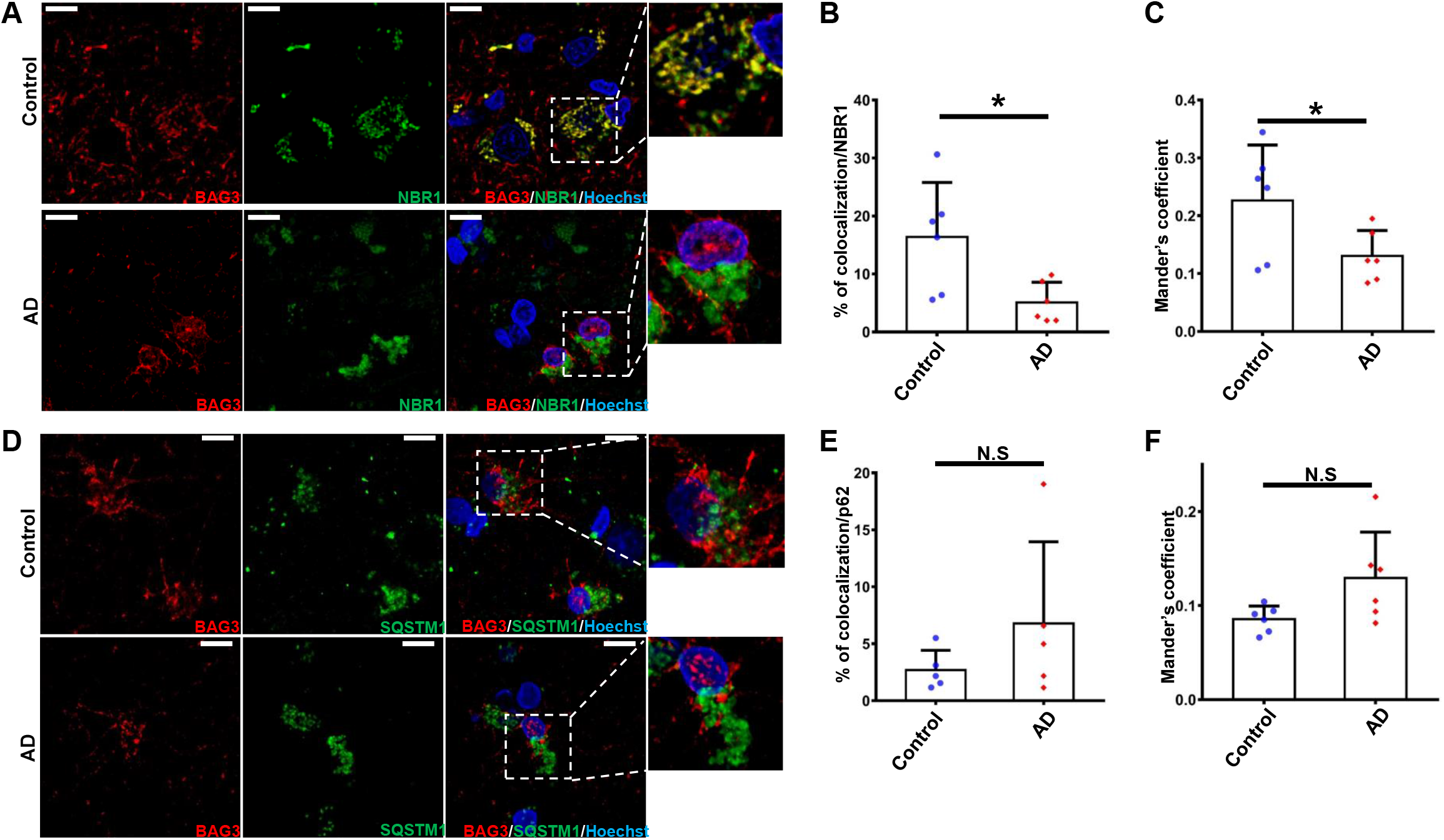
Association of BAG3 and NBR1 is decreased in Alzheimer’s disease brain. (**A**) Representative immunostaining of BAG3 (red) and NBR1 (green) colocalization in human Alzheimer’s disease (AD) brain and age-matched control subjects. Scale bars: 10 μm. (**B**) Quantification of the colocalization between NBR1 and BAG3 based on volume. (**C**) Quantification of co-occurring BAG3 in NBR1, using Mander’s coefficient. (**D**) Representative immunostaining of BAG3 (red) and SQSTM1 (green) colocalization. (**E**) Quantification of the colocalization between SQSTM1 and BAG3 based on volume. (**F**) Quantification of co-occurring BAG3 in SQSTM1, using Mander’s coefficient. Six brains in each group were used for analysis. Data are shown as mean ± SEM using the unpaired Student’s t test; *p < 0.05; **p < 0.01.

**Figure 4.**
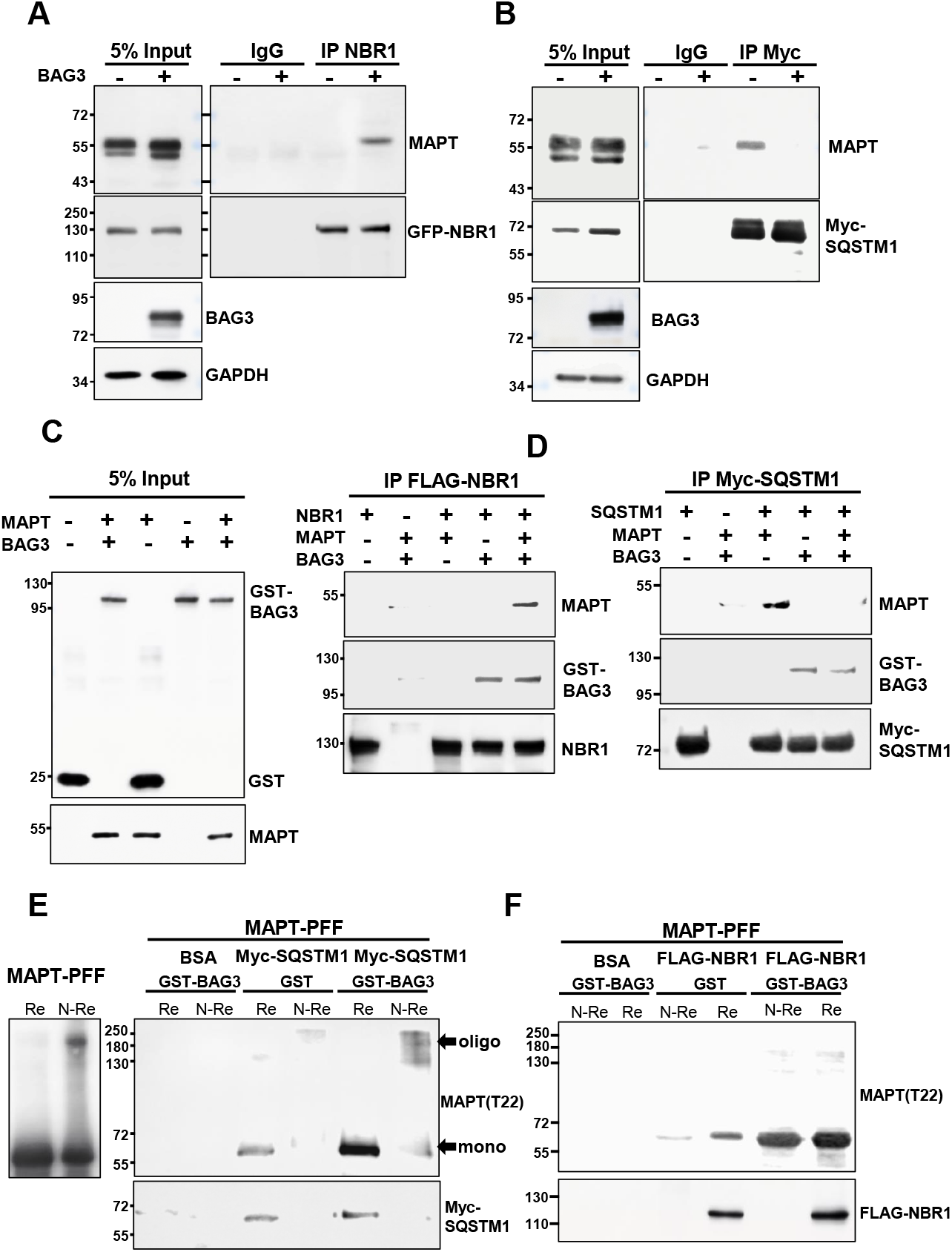
BAG3 regulates the association of autophagy receptors with MAPT. (**A**) BAG3 Null HEK cells were transfected with MAPT, GFP-NBR1 together with BAG3 or control vector. Cell lysates were collected 48 hours post transfection and were immunoprecipitated with NBR1 antibody and immunoblotted with MAPT and NBR1. Five percent of the lysate were used as input control. (**B**) BAG3 Null HEK cells were transfected with MAPT, Myc-SQSTM1 together with BAG3 or control vector. Cell lysates were collected 48 hours post transfection and were immunoprecipitated with Myc antibody and immunoblotted for MAPT and Myc. Five percent of the lysate was used as input control. (**C**) FLAG M2 beads immobilized FLAG-NBR1 or BSA were incubated with purified His-MAPT together with GST or GST-BAG3. The precipitated samples were immunoblotted for His tag, BAG3 and NBR1. Five percent of the His-MAPT together with GST or GST-BAG3 used for pull down of both FLAG-NBR1 and Myc-SQSTM1 were used as input controls. (**D**) Myc beads immobilized Myc-SQSTM1 or BSA were incubated with purified His-MAPT together with GST or GST-BAG3. The precipitated samples were immunoblotted for His tag, BAG3 and Myc. (**E**) Myc beads immobilized Myc-SQSTM1 BSA were incubated with purified pre-formed fibrils together with or without the presence of GST-BAG3. The precipitated samples were eluted with Myc peptide. The eluted sample were split in half in reducing (Re) and non-reducing conditions (N-Re). The samples were immunoblotted with T22 and an antibody to SQSTM1. (**F**) FLAG M2 beads immobilized FLAG-NBR1 or BSA were incubated with purified pre-formed fibrils together with or without the presence of GST-BAG3. The precipitated samples were eluted with FLAG peptide. The eluted sample were split in half in reducing (Re) and non-reducing conditions (N-Re). The samples were immunoblotted for T22 and NBR1. The blots were represented images of three independent experiment.

**Figure 5.**
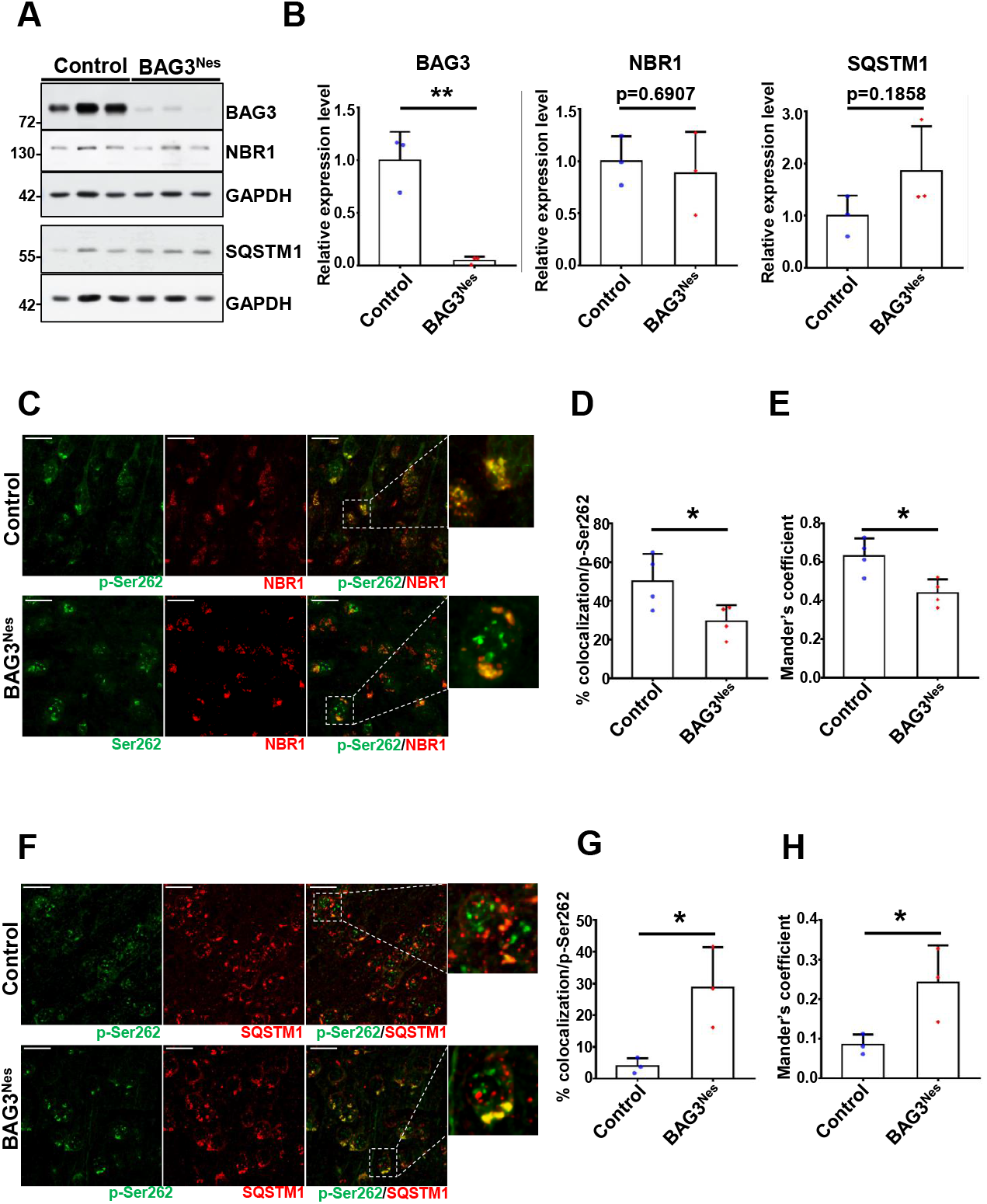
BAG3 regulates the association of p-MAPT with NBR1 or SQSTM1 in mouse brain. Cortices of 24-month-old control and BAG3^Nes^ mice were collected for immunoblotting and immunostaining. (**A**) Cortical lysates from control and BAG3^Nes^ brains were immunoblotted for BAG3, NBR1 and SQSTM1. GAPDH is used as a loading control. (**B**) The graph shows the level of BAG3, NBR1 and SQSTM1 normalized to GAPDH and relative to control. n = 3 for control and BAG3^Nes^. (**C**) Representative immunostaining of p-Ser262 MAPT (green) and NBR1 (red) colocalization. (**D**) Quantification of the colocalization between NBR1 and p-Ser262 MAPT based on volume. (**E**) Quantification of co-occurring NBR1 in p-Ser262 MAPT, using Mander’s coefficient. Three brains in each group were used for analysis. (**F**) Representative immunostaining of p-Ser262 tau (green) and SQSTM1 (red) colocalization. (**G**) Quantification of the colocalization between SQSTM1 and p-Ser262 tau based on volume. (H) Quantification of co-occurring SQSTM1 in p-Ser262 tau, using Mander’s coefficient. Three brains in each group were used for analysis. Scale bar=20um. Data are shown as mean ± SEM using the unpaired Student’s t test, **, p < 0.05.

**Figure 6.**
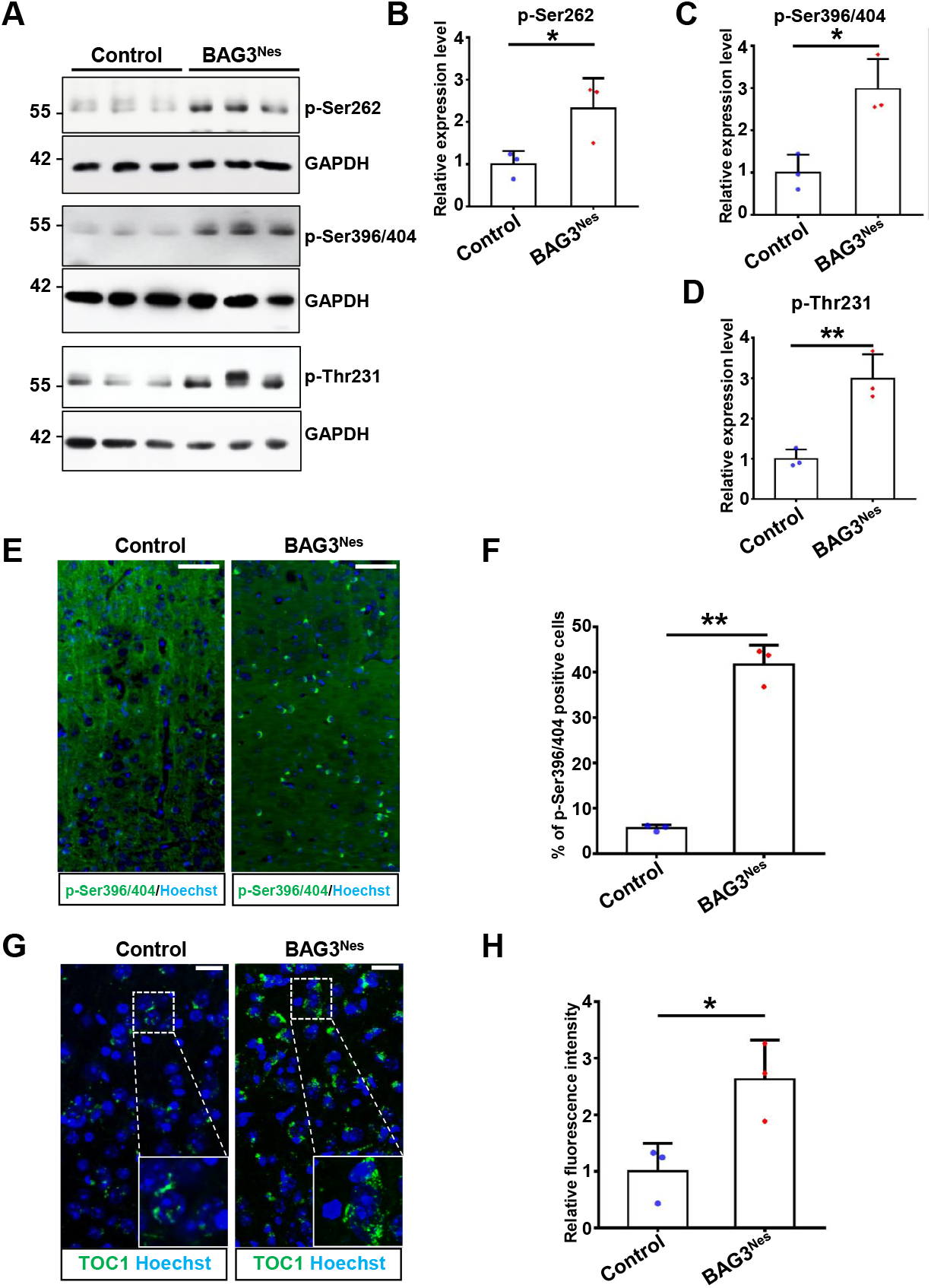
Knock out of BAG3 in mouse brain increased the levels of phosphorylated and oligomeric MAPT. Cortices of 24-month-old control and BAG3^Nes^ mice were collected for immunoblotting and immunostaining. (**A**) Cortical lysates from control and BAG3^Nes^ brains were immunoblotted for phosphorylated MAPT (p-Ser262, p-Ser396/404, and p-Thr231). GAPDH is used as a loading control. (**B-D**) The graph shows the level of phosphorylated MAPT (p-Ser262, p-Ser396/404, and p-Thr231) normalized to GAPDH and relative to control. (**E**) Representative immunostaining of p-Ser396/404 MAPT (green) in the cortex of control and BAG3^Nes^. (**F**) Quantification of the percentage of cells with condensed staining of p-Ser396/404 MAPT. (**G**) Representative immunostaining of TOC1 MAPT (green) in the cortex of control and BAG3^Nes^. (**H**) Quantification of the fluorescence intensity in control and BAG3^Nes^ mice. n = 3 for control and BAG3^Nes^. Scale bars, 20 μm. Data are shown as mean ± SEM. Statistical analysis was performed using the unpaired Student’s t test. *, p < 0.05; **, p<0.01.

### Statistical analysis

All image measurements were obtained from the raw data. GraphPad Prism was used to plot graphs and perform statistical analysis. The statistical tests used are denoted in each figure legend, and statistical significance was defined as *p < 0.05 and **p < 0.01.

## Results

NBR1 and SQSTM1 are autophagy receptors that share similar protein domains but have different targets [28]. To examine the function of NBR1 and SQSTM1 in MAPT clearance, we knocked down NBR1 (Figure 1(A-B)) or SQSTM1 (Figure 1(E-F)) in rat cortical neurons followed by analyses of MAPT levels. Knockdown of NBR1 significantly increased the levels of phosphorylated MAPT including p-Ser262, p-Ser396/404 and p-Thr231 MAPT (Figure 1(C-D)).

In contrast, knockdown of SQSTM1 did not result in changes in phosphorylated MAPT levels (Figure 1(G-H)). The differential impact of the depletion of these two autophagy receptors led us to further examine possible underlying mechanisms.

Our previous findings suggest that BAG3 plays an important role in MAPT clearance in neurons through autophagy [17,18]. Further, it has been reported that BAG3 associates with SQSTM1 in Hela cells [20]. Therefore, we examined the association of BAG3 with NBR1 and SQSTM1. Immunostaining of a control 15-month-old mouse brain revealed that both NBR1 (Figure 2(A)) and SQSTM1 (Figure 2(B)) show a certain degree of colocalization with BAG3 in the CA1 region of hippocampus. However, the colocalization rate of BAG3 with NBR1 was significantly greater than that with SQSTM1, both in pyramidal layer and molecular layer (Figure 2(C)). We further examined the association of BAG3 with NBR1 and SQSTM1 in rat cortical neurons. Rat cortical neurons were transduced with lentivirus expressing scrambled shRNA, or shBAG3 as a negative control. The immunoprecipitation data showed BAG3 co-immunoprecipitated with NBR1 (Figure 2(D)). However, there was less association of BAG3 with SQSTM1 compared to with NBR1 (Figure 2(E)). To extend these findings we examined the colocalization of BAG3 with NBR1 or SQSTM1 in AD brains and the age-matched controls. We found that the association of BAG3 with NBR1 is significantly decreased in AD brains compared with age-matched controls (Figure 3(A-C)). However, there was no significant difference in the co-localization of BAG3 with SQSTM1 between AD brains and the age-matched controls (Figure 3(D-F)). These findings suggest BAG3 differentially associates with NBR1 and SQSTM1 in neurons in mouse and human brains.

These colocalization and association data led us to consider the hypothesis that BAG3 may play a role in directing MAPT recognition by NBR1 or SQSTM1. To test this hypothesis, we expressed MAPT and NBR1 with or without BAG3 in the BAG3 null HEK 293 cells. The immunoprecipitation data showed that NBR1 associated with MAPT in the presence but not in the absence of BAG3 (Figure 4(A)). Interestingly, when the same experiment was carried out with SQSTM1 instead of NBR1, the opposite was true. The SQSTM1 associated with MAPT only in the absence of BAG3 and the expression of BAG3 disrupted this association (Figure 4(B)). To examine if this association was due to direct or indirect binding, we performed FLAG-NBR1 and Myc-SQSTM1 pull-down assays to examine the association of NBR1 or SQSTM1 with BAG3 and MAPT. The FLAG-NBR1 pull-down assay showed that NBR1 directly binds BAG3, and this binding is not affected by MAPT. Further, NBR1 binds MAPT only in the presence of BAG3 (Figure 4(C)). The Myc-SQSTM1 pull-down assay showed that SQSTM1 also directly binds BAG3. However, SQSTM1 only binds MAPT in the absence of BAG3; the presence of BAG3 disrupted the binding of SQSTM1 with MAPT (Figure 4(D)). This in vitro pull-down data is consistent with the findings in HEK cells. These data indicate that BAG3 plays as a regulatory role in modulating the interaction of NBR1 and SQSTM1 with MAPT. Since SQSTM1 has been suggested to preferentially direct the degradation of MAPT aggregates [10], we further examined the effect of BAG3 on the binding of NBR1 and SQSTM1 to MAPT purified pre-formed fibrils (MAPT-PFF). The MAPT-PFFs used in this assay were a mixture of monomeric and oligomeric MAPT (Figure 4(E) left panel). The Myc-SQSTM1 pull-down assay showed that SQSTM1 specifically binds with the oligomeric MAPT, and the presence of BAG3 greatly enhanced the binding of SQSTM1 with oligomeric MAPT (Figure 4(E) right panel, arrow). On the other hand, the FLAG-NBR1 pull-down assay showed that NBR1 specifically binds MAPT in the monomeric state, and the presence of BAG3 greatly enhanced the binding of NBR1 with monomeric MAPT. These findings indicate that BAG3 is acting as a regulator modulating the distribution of different MAPT species to a specific autophagy receptor.

To further examine the role of BAG3 in regulating the association of MAPT with NBR1 and SQSTM1 in vivo, we generated BAG3 conditional knock out mice (Nestin-Cre; BAG3^Fx/Fx^, BAG3^Nes^) in which BAG3 was ablated in neural cells. Western blot data showed that BAG3 levels were significantly reduced in the BAG3 depleted mice (Figure 5(A-B)). The protein levels of NBR1 and SQSTM1 were not significantly different between control and BAG3^Nes^ mice (Figure 5(A-B)). Immunostaining showed that p-Ser262 MAPT exhibited prevalent colocalization with NBR1 in the control mice, while colocalization was significantly decreased in the BAG3^Nes^ mice (Figure 5(C-E)). The opposite was true for SQSTM1. A low level of p-Ser262 MAPT colocalization with SQSTM1 was observed in the control mice, while colocalization was significantly increased in the BAG3^Nes^ mice (Figure 5(F-H)). In addition, we also examined phosphorylated MAPT (p-MAPT) levels in these mice. Immunoblot analysis showed that multiple p-MAPTs including p-Ser262, p-Ser396/404 and p-Thr231 were significantly increased in the BAG3^Nes^ mice (Figure 6(A-D)). Moreover, immunostaining of p-Ser396/404 in control mice showed largely a diffuse staining pattern, while the staining of p-Ser396/404 in the BAG3^Nes^ mice was more punctate and significantly greater than that of the control mice (Figure 6(E-F)). When the brains of the mice were stained with TOC1 antibody which specifically recognizes oligomeric MAPT [23], we found that TOC1 staining was significantly greater in the BAG3^Nes^ mice than that of the control mice (Figure 6(G-H)).

## Discussion

Autophagy is one of the major degradative pathways for aggregated proteins and damaged organelles and therefore, is essential for maintaining neuronal function [29,30]. Selective autophagy, in which select receptors recognize and engage specific cargos, plays a significant role in maintaining protein quality control [31]. Mounting evidence has implicated defective autophagy in the pathogenesis of several major neurodegenerative diseases, particularly AD [32]. In AD it has been postulated that defective autophagy contributes to the pathogenic formation of the key pathological hallmarks of the disease, Aβ plaques and NFTs, which are insoluble aggregates of p-MAPT [33,34]. Further, previous studies have provided evidence that autophagy plays a key role in MAPT clearance [35], and that BAG3 is an important player in autophagy-mediated MAPT clearance [17,18].

In the present study, we found BAG3 can directly bind two autophagy receptors, SQSTM1 and NBR1 in neurons. However, these two autophagy receptors showed dramatically different functions in facilitating MAPT clearance. For example, depletion of NBR1, but not SQSTM1, increased phosphorylated MAPT levels in primary cortical neurons. Further, NBR1-MAPT binding only occurs in the presence of BAG3, while SQSTM1 binding with MAPT only occurs in the absence of BAG3. Moreover, BAG3 promotes SQSTM1 binding with oligomeric MAPT and inhibits its binding with monomeric MAPT; while BAG3 promotes NRB1 binding with the monomeric form of MAPT, which is disrupted in the absence of BAG3. These findings suggest that BAG3 acts as an “instructor” to direct each autophagy receptor to the appropriate cargo for degradation.

Recognition of cargo by selective autophagy receptors is an important component of autophagy pathway [8]. Multiple autophagy receptors such as CALCOCO2, optineurin, and SQSTM1 have been proposed to play a role in regulating the MAPT clearance [36–39]. However, these autophagy receptors showed distinct roles in targeting different MAPT species. For example Xu et al. presented data suggesting that, through autophagy, optineurin regulates the degradation of normal, soluble tau while SQSTM1 primarily regulates the degradation of insoluble MAPT [10]. Directing the autophagy receptors to the different forms of MAPT is a critical step in the degradation of MAPT [10,39]. In our present study, we found that the presence of BAG3 directs monomeric MAPT to NBR1 and not SQSTM1 in either HEK cells or an in vitro pull-down assay. This is consistent with a previous finding in which NBR1 was found to target a soluble cargo through ESCRT pathway for degradation [11]. Since BAG3 also regulates MAPT degradation through the ESCRT pathway, it can be speculated that the promotion of NBR1-MAPT interactions by BAG3 may also facilitate MAPT clearance by the ESCRT pathway [19,40]. Interestingly, removal of BAG3 disrupted the binding of monomeric MAPT to NBR1 but directed monomeric MAPT to SQSTM1. As SQSTM1 selectively regulates the degradation of oligomeric MAPT species [10], the monomeric MAPT that binds to SQSTM1 may not be properly degraded. This may also explain our observation that neural cell specific depletion of BAG3 enhanced the association of p-MAPT with SQSTM1 but also resulted in the accumulation of p-MAPT. Further, when NBR1 was incubated with a mixture of monomeric and oligomeric MAPT, it preferentially bound monomeric MAPT, in contrast to SQSTM1 which preferentially bound MAPT in the oligomeric state. Most importantly, the presence of BAG3 promoted the binding of NBR1 with monomeric MAPT as well as the binding of SQSTM1 with oligomeric MAPT. Taken as a whole, these data clearly demonstrate that NBR1 and SQSTM1 selectively bind different states of MAPT.

Our findings indicate that BAG3 plays a novel role in directing the interaction of NBR1 and SQSTM1 with MAPT with a specific form of tau to mediate its clearance. The fact that BAG3 mediates the form of MAPT selectively recognized by SQSTM1 or NBR1 is consistent with our findings in neurons and the mouse model. Monomeric MAPT is the predominant form in primary neurons without oligomerization induction [41,42]. Therefore, the finding that knockdown of NBR1, but not SQSTM1, lead to an increase in p-MAPT is expected as NBR1 preferentially interacts with monomeric MAPT in the presence of BAG3. The decreased association of BAG3 with NBR1 that we observed in AD brains as well as in BAG3^Nes^ mice may also contribute to the MAPT accumulation.

In summary, we have provided direct evidence of the district roles of NBR1 and SQSTM1 in binding with monomeric and oligomeric MAPT, respectively, to promote their clearance. More importantly, we also found that BAG3 regulates the specificity of the recognition of monomeric MAPT by NBR1 and oligomeric MAPT by SQSTM1. Knockout of BAG3 disrupted this specificity and lead to increased accumulation of p-MAPT and oligomeric-MAPT in mouse brain. Overall, the results of this study indicate that BAG3 is a key regulator of MAPT clearance mechanisms and thus for maintaining MAPT proteostasis in neurons.

## Abbreviations

AD: Alzheimer’s disease
BAG3: BCL-2-associated athanogene 3
BSA: Bovine serum albumin
CERAD: Consortium to Establish a Registry for Alzheimer’s Disease
ESCRT: endosomal sorting complexes required for transport
GST: Glutathione S-transferases
MAPT: microtubule-associated protein tau
NBR1: Neighbor of BRCA1 gene 1
NFT: neurofibrillary tangles
PMI: post-mortem interval
SQSTM1: Sequestosome-1

## Acknowledgements

This work was supported by the National Institutes of Health (NIH) (R01 AG073121. [to GVWJ]) and the Alzheimer foundation (AARF GR531565 [to HL]).

GJ contributed to the study conception and design, edited the manuscript, and provided funding. HL contributed to the study conception and design, performed experiments, analyzed the data, interpreted the experiments, wrote the manuscript, and provided funding. CD and SS contributed to the study conception and design, performed experiments, analyzed the data. DK performed experiments, analyzed the data.

We thank Dr. C. Pröschel, University of Rochester, for providing us with the pHUUG and FigB vectors; Dr. P. Davies for the gift of phospho-MAPT (Ser396/404) (PHF1) antibodies; and Dr. P. Dolan for the gift of phospho-MAPT (Ser262) antibody. Dr. N. Kanaan, Michigan State University for the gift of TOC1 antibody. T. Johansen, University of Tromso for Myc-SQSTM1 plasmid. Dr. Jeff Kuret, Ohio State University for pT7-htau40 vector. Brain samples were obtained from the University of Florida Neuromedicine Human Brain and Tissue Bank (UF HBTB) with informed consent of the patients or their relatives and the approval of the local institutional review boards. The UF HBTB is supported by the Florida Alzheimer’s Disease Research Center (P30AG066506). We would also thank Sudarshan Ramanan for assistance with some of the immunoblots.

## Figure legends

**Supplementary Table 1.**
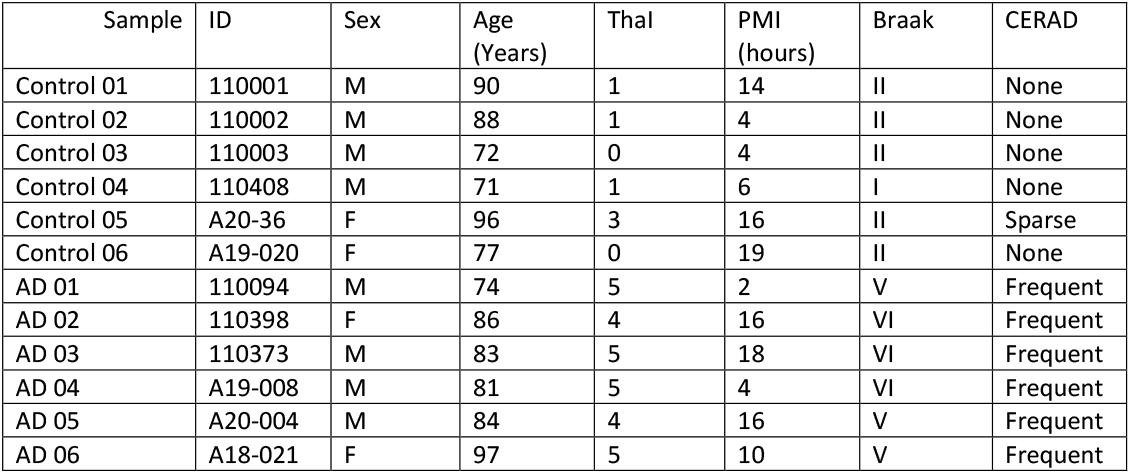

